# The Computational Analysis of *Plasmodium falciparum* Heat Shock Proteins Reveals an Interplay with Polyamines

**DOI:** 10.1101/2023.05.11.540332

**Authors:** Godlo Sesethu, Maxam Nombalentle, Mthembu Yamnkela, Mpumza Anelisa, Stanley Makumire, Noxolo Mkwetshana, Krishna K Govender, Xolani H Makhoba

## Abstract

The current drugs available in the market are not effective due to growing numbers of resistance to the causative agent of malaria. There are various *Plasmodium* parasites, of which *Plasmodium falciparum* is the main cause of morbidity and mortality reported worldwide. Therefore, there is an urgent need to come up with an innovative and effective treatment for this disease. Polyamines play a major role in the parasite’s well-being and growth, while heat shock proteins keep the proteomics of the parasite in good shape. In this study, *In Silico* analysis of the interaction between putrescine, spermidine, spermine, and heat shock proteins was carried out to establish the binding site for drug discovery. Computational tools such as Bioedit, PROCHECK, KNIME Hub, and Schrodinger were used. The results revealed interactions between polyamines and heat shock proteins with glutamine and aspartic acid being common amino acids where interaction occurs between the chaperones and polyamines. MD shows a strong interaction between PfHsp70-1 and putrescine, but the best interaction is observed for PfHsp70-1 and spermidine. Based on these results, a follow-up study will be conducted to establish the synthesis of drugs that will be used as targets for both polyamines and heat shock proteins to eradicate malaria.

**Author’s summary:** The emanation and spread of Plasmodium parasites that are resistant to antimalarial therapy is one of the main problems in the treatment of malaria. This is a result of the Plasmodium parasite’s ongoing evolution and the creation of novel strategies for surviving drug toxicity. Studies of antimalarial drug development have been focused on polyamine biosynthesis by targeting precursors such as ornithine decarboxylase, adenosylmethionine decarboxylase, and spermidine synthase and protein-protein interactions between *Plasmodium falciparum* chaperones spotting out Hsp90, Hsp70, and Hsp40 as potential targets with little attention being paid to the interaction between polyamines and molecular chaperones. Therefore, to study these interactions the binding sites of all 3D structures were identified using SiteMap, and a docking was performed using the Schrödinger software with OPLS4 force field and XP.

## 1. Introduction

Malaria is a mosquito-borne disease, it is one of the most predominant parasitic diseases that threaten human life. Malaria affects about half of the world’s population. Most deaths occur in Sub-Saharan Africa and high-risk areas such as Southeast Asia, the Eastern Mediterranean, the Western Pacific, and the Americas [1]. In 2019, the World Health Organization reported approximately 229 million cases and 409 000 malarial deaths. Sub-Saharan Africa accounted for 94% of these deaths [2]. This parasitic disease is transmitted to humans by female *Anopheles* mosquitoes. Of the more than 430 *Anopheles* species known to exist, approximately 40 can transmit malaria. In humans, five *Plasmodium* species are causative agents of malaria infection: *Plasmodium falciparum, Plasmodium vivax, Plasmodium ovale, Plasmodium malariae*, and *Plasmodium knowlesi*.

*Plasmodium falciparum* is the most common malaria parasite as it accounts for over 99% of malaria cases in Africa, 71,9% in the West Pacific region, 69% in the Mediterranean, and 62, 8% in South-East Asia [2]. The most vulnerable groups at risk of contracting malaria include pregnant women and children under the age of 5, with 67% of malaria death cases being children under the age of 5 [1]. Malaria mostly affects developing tropical and subtropical countries. In these countries, outcomes of malaria include loss of education where children miss school, and adults not being able to provide for their families due to physical and intellectual distress caused by cerebral damage. The disease results in financial burdens for both the government and individuals. WHO reported that in Africa, direct costs of malaria sum up to approximately US$12 million annually, while households spend a quarter of their income on treating malaria at home.

The drug of choice for treating malaria is determined by several variables including age, the severity of the infection, baseline immunity, parasite sensitivity, and drug cost and availability. In the early 1900s, quinine and its derivatives were widely used to treat malaria. Many of these anti-malarial drugs are no longer in use due to *P. falciparum* and *P. vivax* resistance against the drugs and undesirable side effects from compounds such as mepacrine. However, some quinine derivatives such as mefloquine are still in use in combination with artesunate as binary combination drugs [4]. Currently, classes of drugs used for malaria include artemisinin, antifolates, 4-aminoquinolines, 8-aminoquinolines aryl-amino alcohols, and antibiotics. Artemisinin is the most widely used antimalarial drug globally due to its efficacy against all forms of *P. falciparum* strains by inhibiting the parasite’s phosphatidylinositol-3-kinase during the early ring stage [5] with artemisinin combination therapies (ACTs) being recommended to treat both complicated and uncomplicated *P. falciparum* malaria. However, gene modification in the *Plasmodium* parasite results in resistance of the parasite to antimalaria drugs as the genes that encode target regions mutate, and this raises a crucial need for new innovative, and effective antimalarial drugs [8]. Newly developed drugs should at least be able to terminate the parasite growth at any stage, in the erythrocytic stage, mosquito stage, and human liver stage [9] as a way of changing the usual sequence of drug targets.

Positively charged organic compounds with two or more amino groups found in cells of living organisms are referred to as polyamines [10], and they play an important part in cell division and differentiation of *P. falciparum* [10,11]. These include diamine, putrescine, and triamine spermidine as well as tetramine spermine, which are synthesized uniquely by the action of S-Adenosylmethionine decarboxylase (AdometDC) and ornithine decarboxylase (ODC) functioning as a single enzyme, and spermidine synthase. The absence of spermine synthase, in *P. falciparum*, makes spermidine synthase responsible for both spermidine and spermine synthesis [11].

The malaria parasite survives between the mosquito host and human host. During transmission between these two hosts, the parasite moves from a poikilothermic mosquito to a warm-blooded human, causing severe heat shock to the parasite. Thus, heat shock proteins play a vital role in the adaptation survival of the parasite inside the human host [3]. Heat shock proteins, which are a part of a large family of molecular chaperones, are well-known for their function in the re-folding, maturation, and degradation of proteins [13]. These are divided into small heat shock proteins and major heat shock proteins [12,13], of which the major ones have a molecular mass of greater than 43kDa, and small heat shock proteins have molecular weights of less than 43kDa [13]. Small heat shock proteins with molecular weights of 15-43kDa are called heat shock protein β and are particularly known for their role in protecting cells from stress [14].

## 2. Results and discussion

### 2.1 Sequence retrieval and Genomic Analysis

Sequences were retrieved from the NCBI database (https://www.ncbi.nlm.nih.gov/). Genomic analysis showed that HSP20 and HSP 70 are located on the same chromosome (chromosome 8), while HSP40 is located on chromosome 13, HSP60 on chromosome 10, and HSP90 on chromosome 7 (Supplementary Table 1). The bioinformatics analysis of *P. falciparum* 3D7 heat shock proteins showed that these major HSPs have a few exons and introns in them, with HSP40 and HSP70 having 1 exon and no introns, HSP60 and HSP 90 have 2 exons and 1 intron, and HSP 20 has 3 exons and 1 intron.

### 2.2 Multiple sequence alignments of Pf Hsps and their homologs

All the heat shock proteins were aligned with *Saccharomyces cerevisiae, E. coli*, and humans in BioEdit, using ClustalW and BLOSSUM62 matrix. The similarity index between *P. falciparum and E. coli* was HSP20, HSP40, HSP60, HSP70 and HSP 90 at 20,63%, 19,34%, 67,41%, 58,30%, and 54,15% respectively. While the similarity index between *P. falciparum* and *S. cerevisiae* HSP40, HSP60, HSP70, and HSP90 was higher at 32,90%, 71,92%, 79,32%, and, 75,7% respectively except for HSP20 which was 12,61%. This is due to *S. cerevisiae’s* small HSP (HSP20 family) being present as HSP26, a 26 kDa protein instead of 20 kDa.

### 2.3 Homology modelling and structure validation

All PDB structures (Figure 1) were predicted with Phyre2, analyzed and confirmed with PROCHECK on PDBSum, and visualized with PyMOL v2.x. The projected structure of PfHSP20 comprises 2 sheets, 4 beta hairpins, 7 strands, 6 helices, 15 beta turns, and 6 gamma turns, according to motif analysis. This demonstrated that the created structure was suitable for further investigation. The 3D structure of PfHSP40 reveals that the protein has 5 helices, 9 helix-helix interactions, and 2 beta twists. The Ramachandran plot revealed that all of the residues were in core areas. As a result, the structure merits additional investigation. The 3D structure of HSP60 revealed 5 sheets, 2 beta alpha beta units, 3 beta hairpins, 1 psi loop, 2 beta bulges, 17 strands, 27 helices, 36 helix-helix interactions, 25 beta turns, and 5 gamma turns. According to the Ramachandran plot data, this structure included 95.8 percent of the residues in the most desired areas, 3.9 percent of the residues in allowed regions, 0.4 percent of the residues in the generously allowed regions, and none in the banned regions. The motif description of PfHSP70’s 3D structure is the same as that of PfHSP60. The plot of PfHSP70 Ramachandran indicated that 92.0 percent of the amino acid residues are in the core areas, 7.3 percent are in allowed regions, 0.5 percent are in generously allowed regions, and 0.2 percent are in banned regions. The generated PfHSP90 structure has 1 sheet, 4 beta hairpins, 2 beta bulges, 9 strands, 9 helices, 15 helix-helix interacts, 13 beta turns, and 2 gamma turns, with the Ramachandran plot indicating that 92.4 percent of the residues are in core regions and 7.6 percent are in allowed regions.

**Figure 1:**
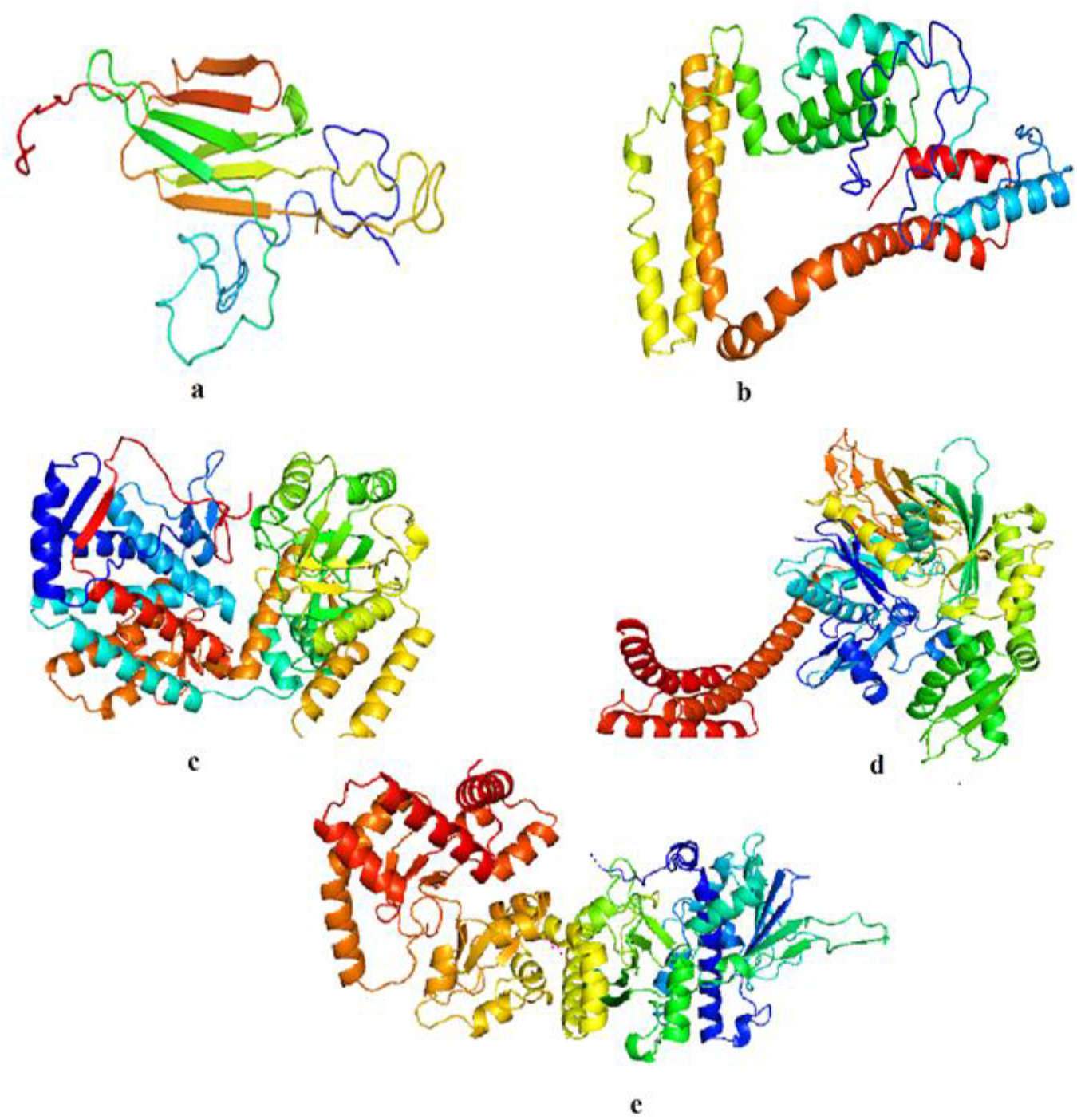
Three dimensional structures of P. falciparum 3D7 (a)HSP20, (b)HSP40, (c)HSP60, (d)HSP70, and (e)HSP90 in cartoon format as visualised on PyMol

### 2.4 Docking sites identified in the heat shock proteins

The heat shock proteins’ 3D models were modelled using Phyre2 [6]and these were used to identify possible binding sites. . Active sites were identified by making use of the Sitemap tool [Must reference] provided within the Schrodinger software suite. For a site to be considered a good potential docking site the following needs to hold true, sitescore > 1, dscore > 1 and a volume > 225. Based on the results obtained (Perhaps we can put the table of data into supplementary) those sites provided in Figure 2 where the ones chosen for docking as they obeyed the criterion specified. This approach was chosen as opposed to blind docking since it was computationally more appealing and has been shown to provide good estimates for active site identification. [Halgren, T., “Identifying and Characterizing Binding Sites and Assessing Druggability,” J. Chem. Inf. Model., 2009, 49, 377–389. Halgren, T., “New Method for Fast and Accurate Binding-site Identification and Analysis,” Chem. Biol. Drug Des., 2007, 69, 146–148.]

**Figure 2.**
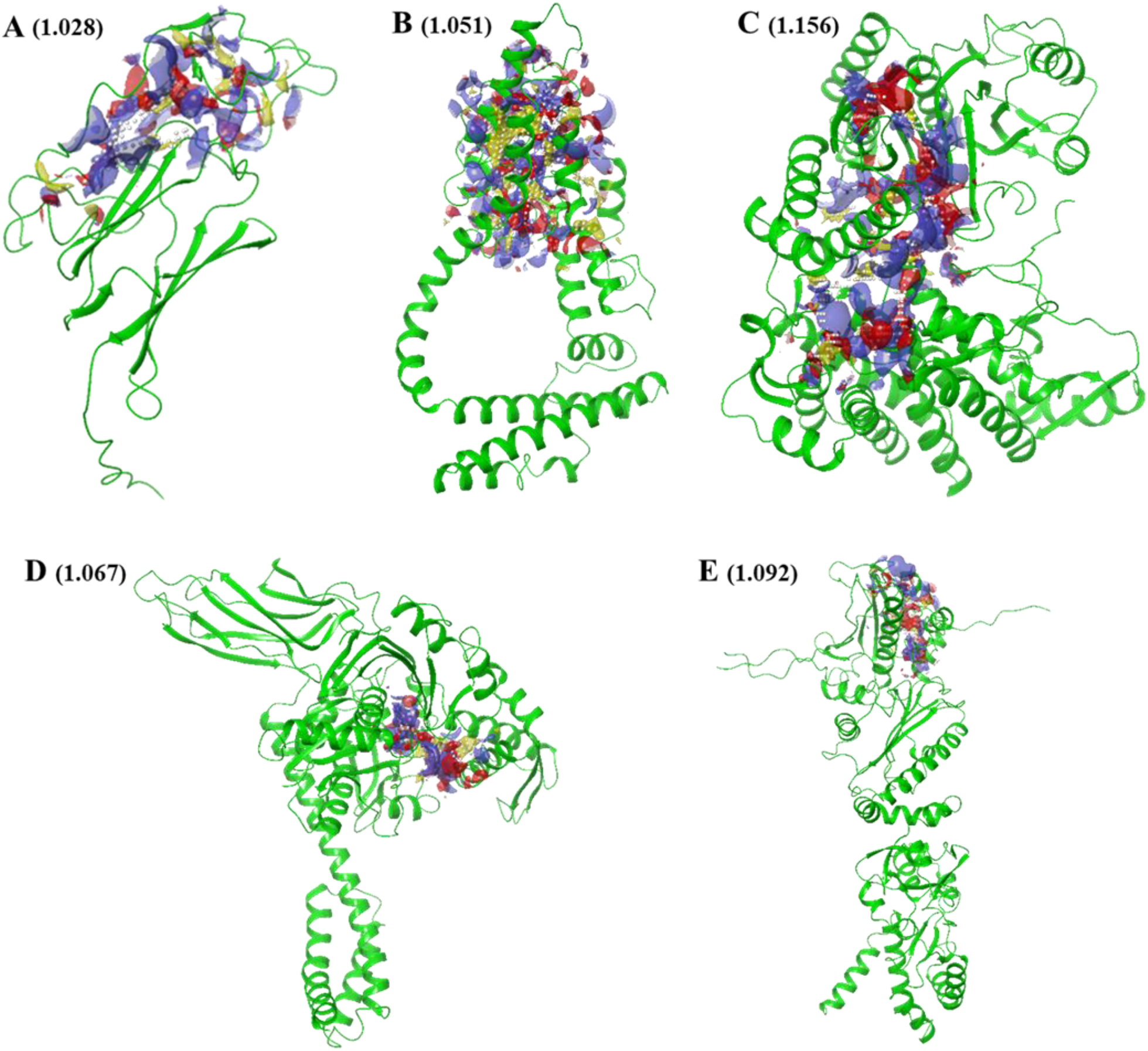
Active site identification of (A) HSP20, (B) HSP40, (C) HSP60, (D) HSP70 and (E) HSP90 identified via SiteMap. Site scores are given in brackets.

### 2.5 Molecular docking

Those ligand conformations that provided the lowest docking scores after being bound to the active sites of the heat shock proteins are provided in Table 1.

**Table 1.** Docking scores of Putrescine, Spermidine, and Spermine bound to the active site of heat shock proteins

| Ligand Name | CID | HSP20 | HSP40 | HSP60 | HSP70 | HSP90 |
| --- | --- | --- | --- | --- | --- | --- |
| Putrescine | 3452892 | -3.19 | -5.41 | -5.95 | -6.63 | -6.68 |
| Spermidine | 6992097 | -4.54 | -8.18 | -7.34 | -7.99 | -6.19 |
| Spermine | 446718 | -3.07 | -5.86 | -8.66 | -4.41 | -4.76 |

In the case of Put, the docking scores appear to drop as the protein structure increases in size. This could be due to the binding pocket getting larger as the proteins get bigger and since Put is a small ligand it can fit better within the active site. For Spd the lowest docking score (−8.18 kcal/mol) is obtained when the ligand is bound to HSP40, while for Spn the lowest docking score (−8.66 kcal/mol) is obtained for HSP60. Figures 3 – 5 provide the ligand interaction diagrams for the different ligands bound to the active sites of the heat shock proteins considered in this work.

**Figure 3.**
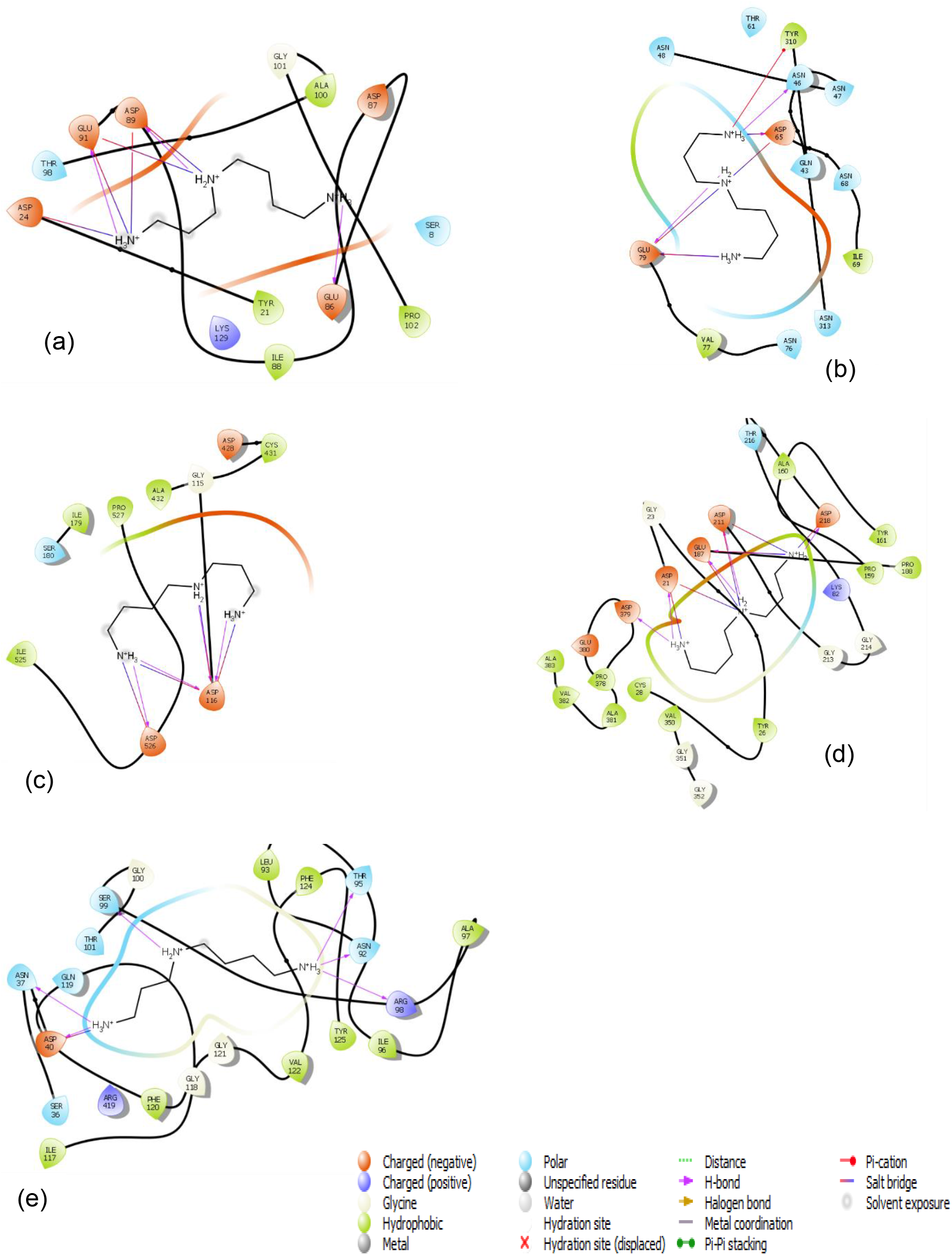
Ligand interaction diagram of (a) HSP20, (b) HSP40, (c) HSP60, (d) HSP70 and (e) HSP90 bound to Put.

A summary of all the hydrogen bond interactions and salt bridges formed between the ligand and amino acid residues contained within the active site of the proteins is provided in Table 2. For HSP20 the amino acid Asp89 has a hydrogen bond that takes place between all three ligands, while Glu91 has a hydrogen bond present for ligands Put and Spd. Asp24 shows a hydrogen bond between Spd and Spn. It is this combination of salt bridges and hydrogen bond interactions that adds to the protein-ligand complex stability.

**Figure 4.**
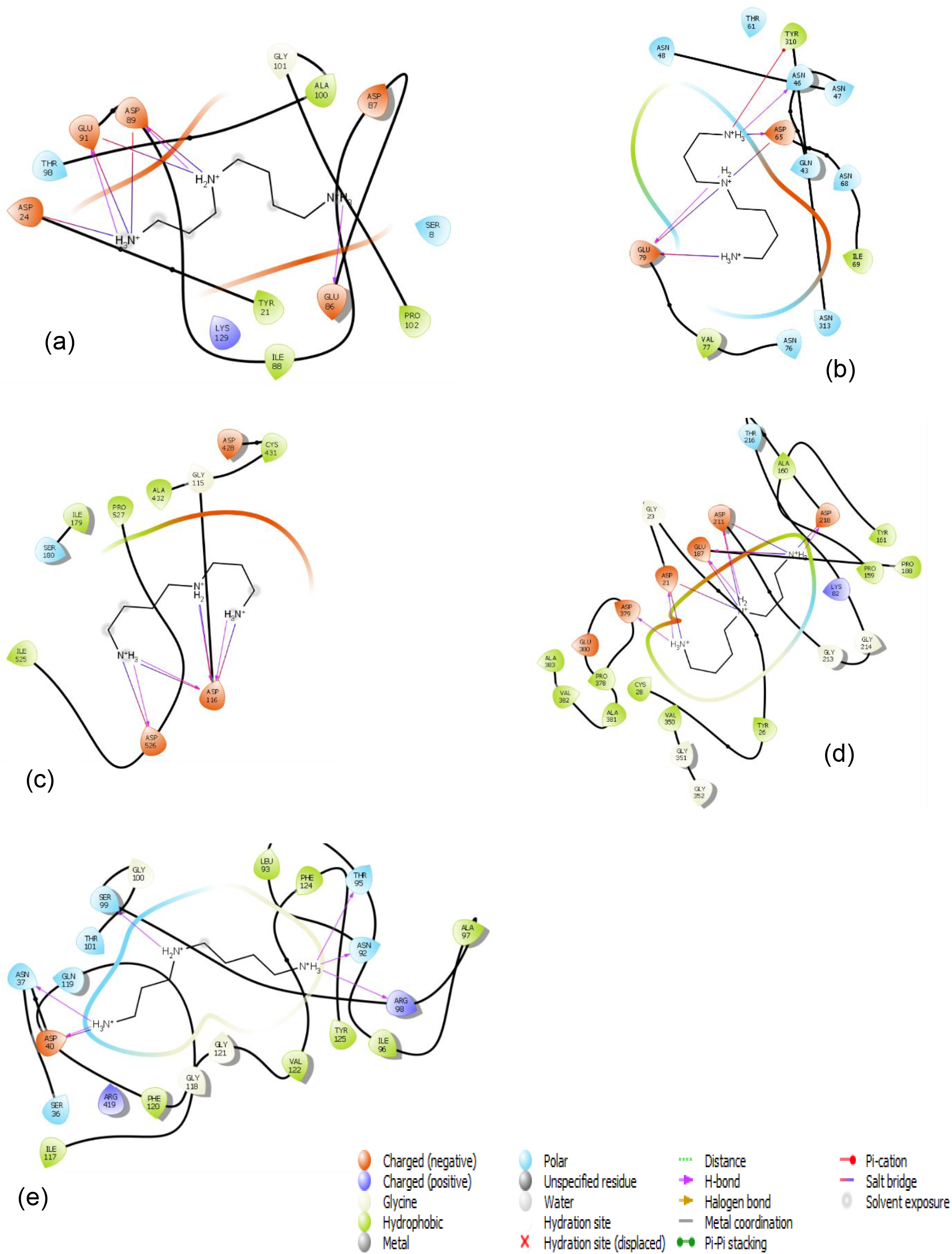
Ligand interaction diagram of (a) HSP20, (b) HSP40, (c) HSP60, (d) HSP70 and (e) HSP90 bound to Spd.

**Figure 5.**
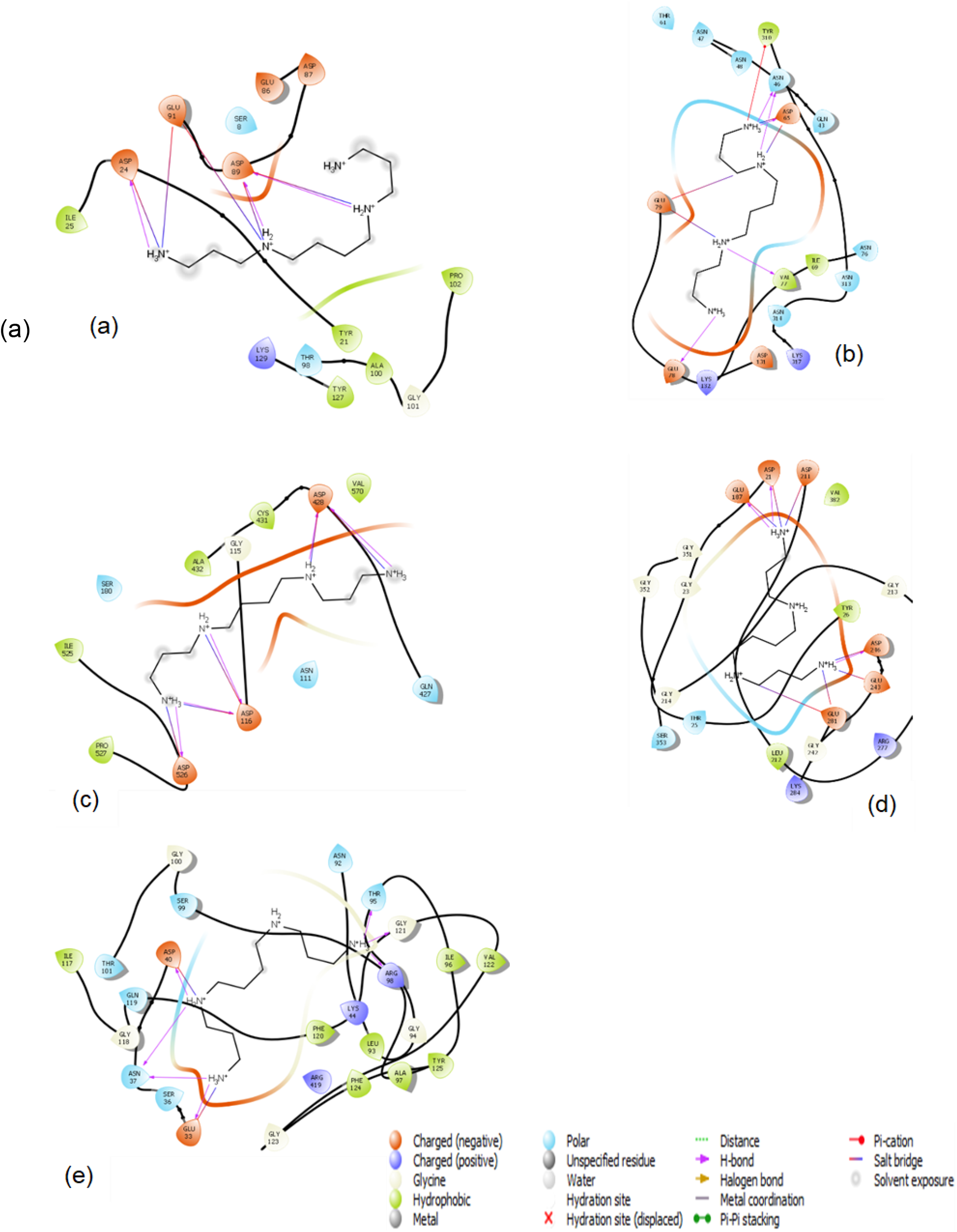
Ligand interaction diagram of (a) HSP20, (b) HSP40, (c) HSP60, (d) HSP70 and (e) HSP90 bound to Spn.

**Table 2.**
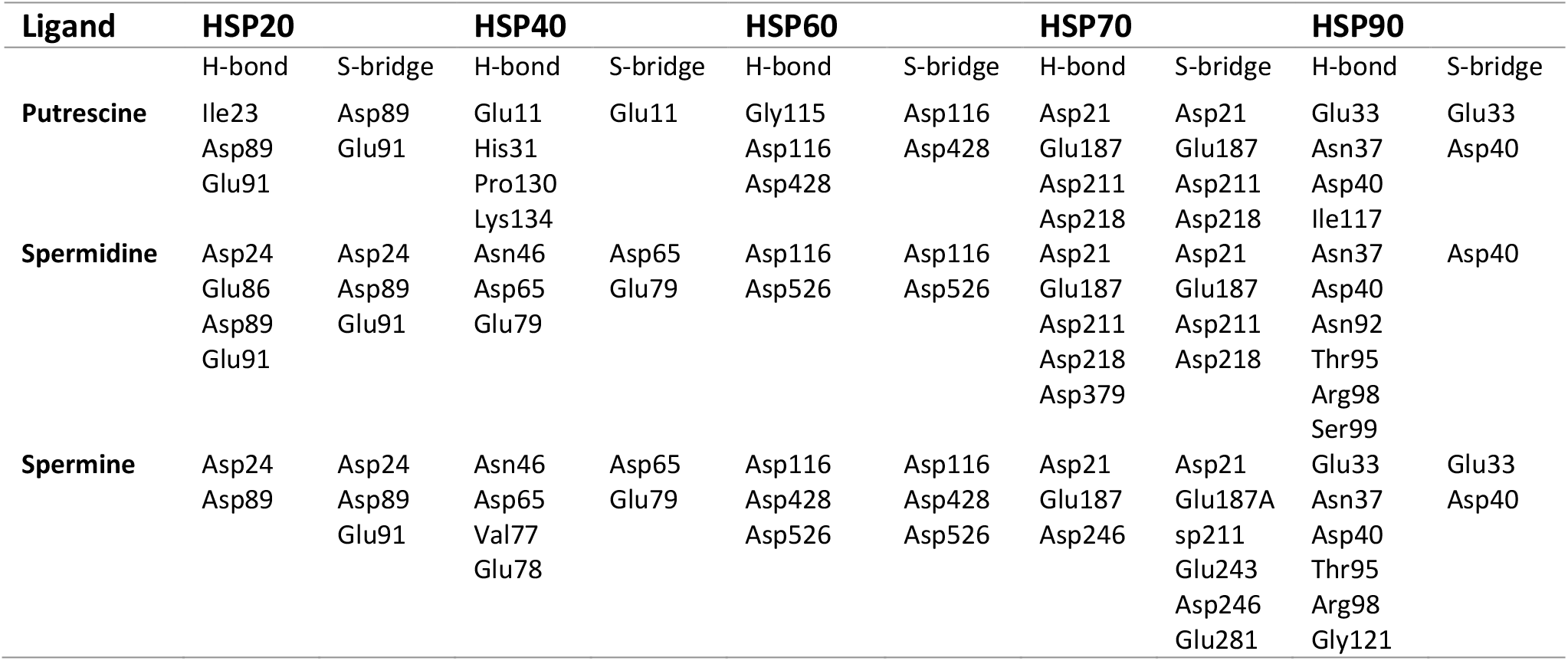
Amino acid residue interactions obtained from the ligand interaction diagrams

### 2.6 Molecular Dynamics

Before we investigated the results obtained for the various HSP complexes we first needed to make sure that the environment chosen for our MD studies was appropriate. Figure 6 below shows the root mean square deviation (RMSD) results obtained for 200 ns simulations of (a) HSP20, (b) HSP40, (c) HSP60, (d) HSP70, and (e) HSP90 when there were no ligands present within the active site of the proteins.

**Figure 6.**
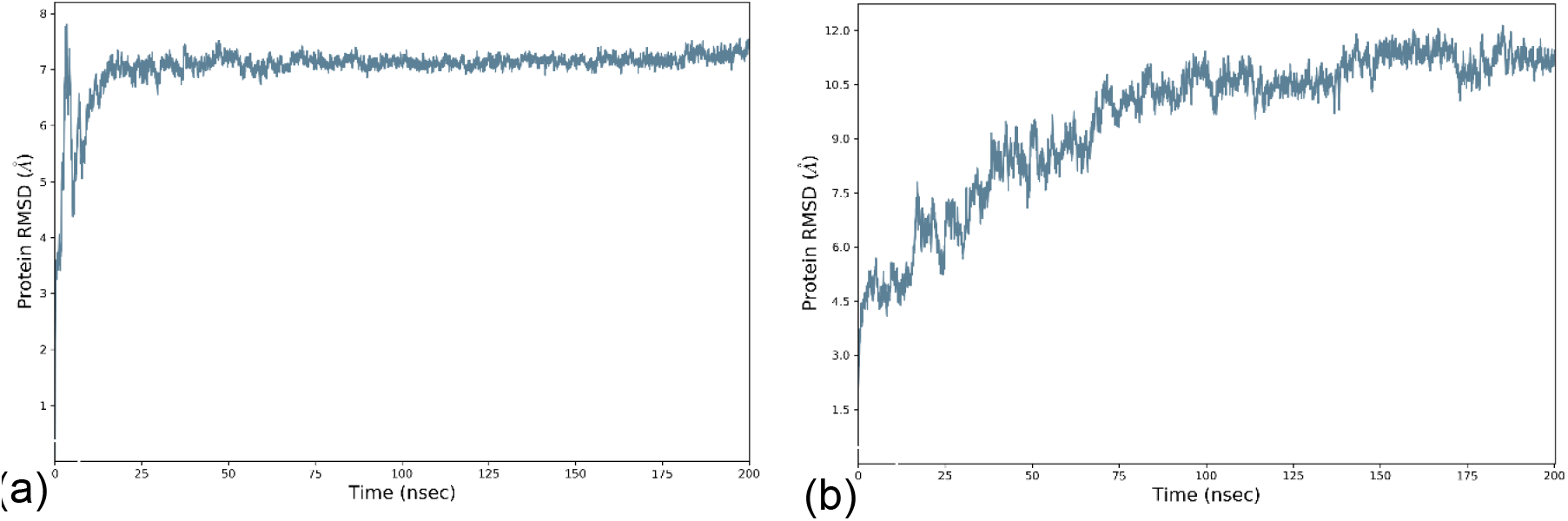
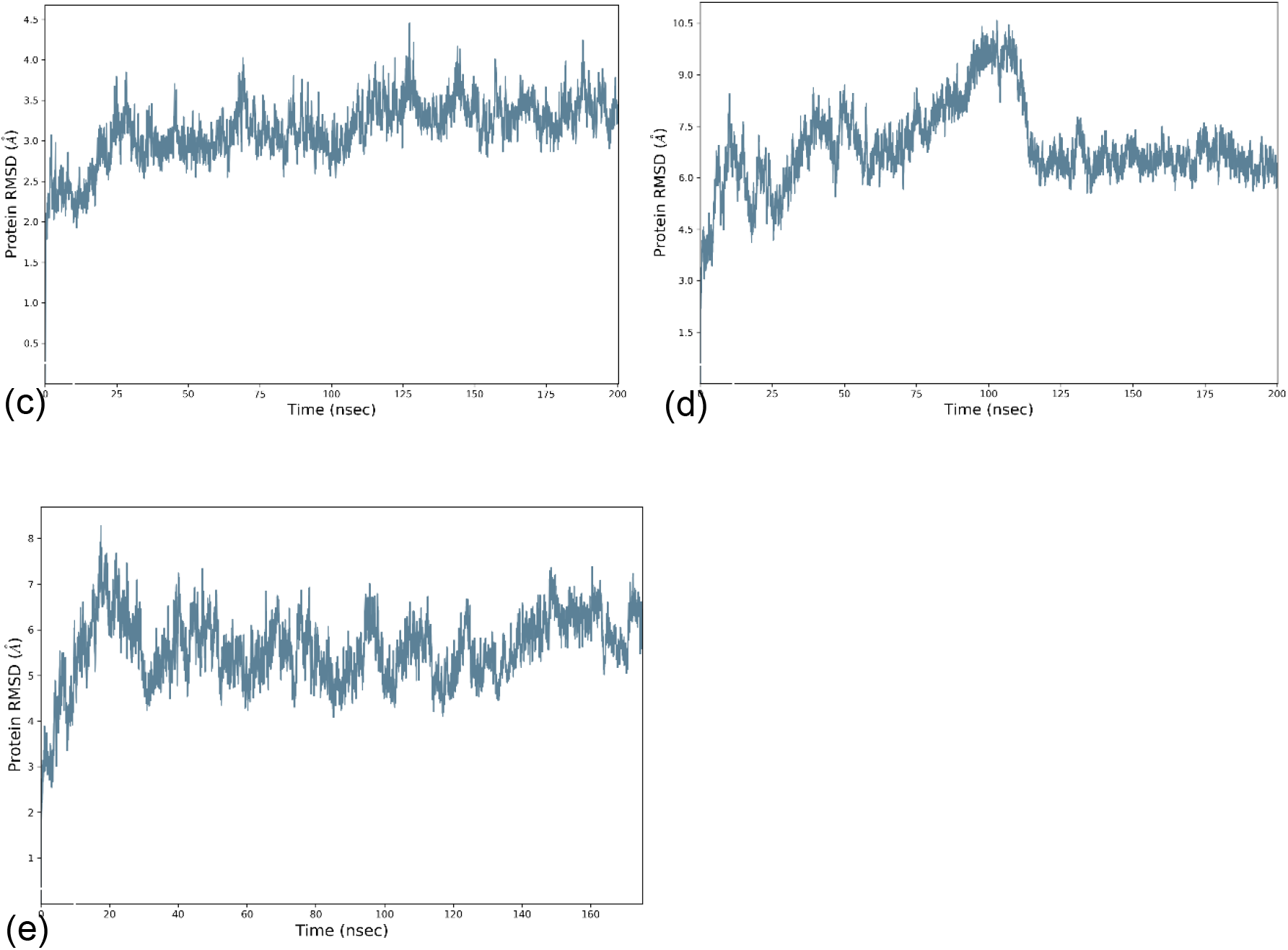
RMSD of alpha carbon backbone for free (a) PfHsp20sp, (b) PfHsp40, (c) PfHsp60, (d) PfHsp70 and (e) PfHsp90 during 200 ns MD simulation.

The RMSD is used for measuring the changes that take place to the protein while present in a solvated environment [23]. A protein’s stability can be determined by the deviations produced during the MD simulation. Figure 6 does show that PfHsp20, PfHsp40, and PfHsp70 stabilize within the 200 ns, but PfHsp60 and PfHsp90 do still seem to have some excess fluctuations towards the end of the 200 ns. Although this was the case it was decided that we will make use of a maximum of 200 ns for the protein-ligand complex simulations due to the limited resources that we had access to at the high-performance computing centre.

Molecular dynamics simulations of the Hsps with the polyamines showed that Putrescine best binds to PfHsp60 and PfHsp70 (Figure 7) compared to the other Hsps. Put seems to remain strongly bound to the active site of PfHsp60 for around 125 ns before the ligand begins to drift away from the active site, reaching distances of up to 14 Å from the starting structure (Figure 7A). For ligands to be considered good inhibitors, they need to remain bound to the active site for longer periods. From the RMSD provided in Figure 7A, we can see that a conformation change in the protein structure results in the ligand leaving the protein’s active site. However, for PfHsp70 (Figure 7B) the ligand Put does have the potential to be a ligand that can bind to the protein for the entire 200 ns timeframe as we see the highest deviation that the ligand has from its starting structure is about 6.0 Å, which takes place around 100 – 120 ns. With that said 6.0 Å is not exactly a small deviation from the active site so this might be a weak binding ligand should it possess the ability to bind to the active site. As was the case for PfHsp60, changes in the conformation of PfHsp70 are what bring about the changes found in the ligand, when the protein RMSD goes up then so too does the ligand RMSD and vice versa.

**Figure 7.**
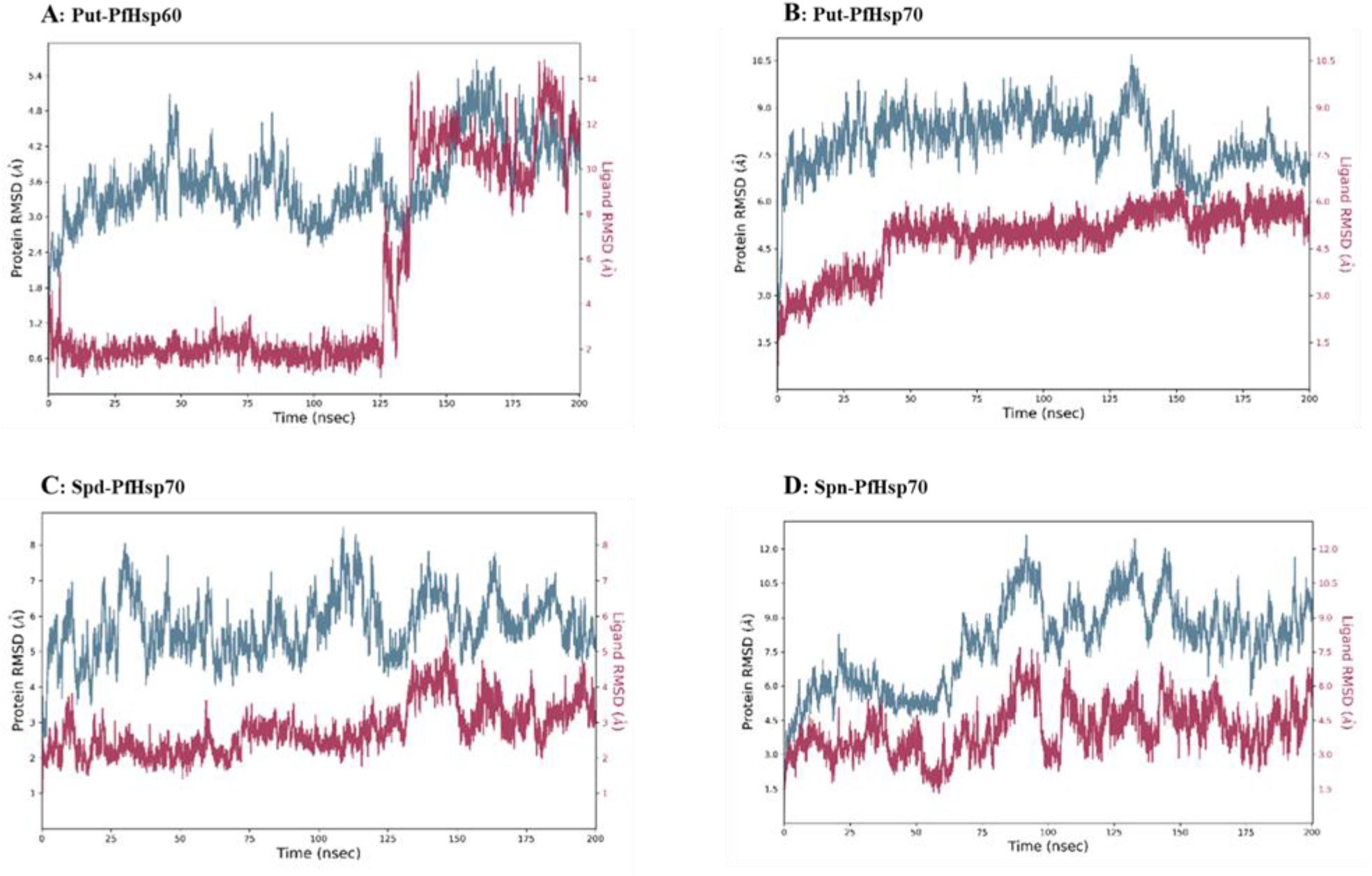
RMSD of the alpha carbon backbone and ligand fluctuations for (A) Put-PfHsp60, (B) Put-PfHsp70, (C) Spd-PfHSP70, and (d) Sp-PfHSP70 during 200 ns MD simulation.

Spd and Spn were only observed to stay bound to PfHsp70 for longer compared to the other Hsps tested. For PfHsp70-Spd (Figure 7C) the ligand does appear to be bound to the active site of the protein for the duration of the MD simulation with the largest difference for the ligand being 5.5 Å from its starting structure. When compared to the PfHsp70-Put complex (difference of 6 Å) it does seem to indicate that Spd is a better binder when compared to Put. The PfHsp70-Spn (Figure 7D) seems to show weak binding as the ligand fluctuations appear to reach up to 7.5 Å in this case.

## 3. Materials and Methods

Appendix 1 contains a list of online web servers used for bioinformatic analyses used in this study.

### 3.1 Sequence retrieval

Sequences of selected heat shock proteins from *Plasmodium falciparum* 3D7 were retrieved from the PlasmoDB database. This database provides substantial Plasmodium spp. genome, proteome, and metabolome information. Genomic characteristics of the selected heat shock proteins were obtained from the National Centre of Biotechnological Information (NCBI) database by selecting the gene ID of the selected HSP to identify the protein’s gene and chromosomal location.

### 3.2 Multiple sequence alignment

Protein sequences of Hsp20, Hsp40, Hsp 60, Hsp70, and Hsp90 from *Escherichia coli, and Saccharomyces cerevisiae* were retrieved from the NCBI. The sequences, together with the *P. falciparum* sequences were aligned using the Bio-edit tool.

### 3.3 Homology modelling and structure validation

Phyre2, a protein structure prediction database was used. to generate 3D model structures using sequences of the *P. falciparum* 3D7 selected HSPs. The modeled structures were visualized with PyMol. The structures were then validated using PROCHECK on the online tool PDBSum.

### 3.4 Protein preparation for Docking and Molecular Dynamics

All the heat shock proteins (PfHsp20, PfHsp40, PfHsp60, PfHsp70 and PfHsp90) obtained from homology modelling were prepared with the aid of the protein preparation wizard provided in Schrodinger 2022-1 [18]

### 3.5 Site Mapping

Since the proteins were obtained via homology modelling there were no active sites to work from. Instead of making use of a blind docking approach, we made use of the SiteMap [19] the tool provided in Schrodinger 2022-1. The tool allows for the identification of binding sites whose size, functionality, and extent of solvent exposure meet certain criteria. The OPLS4 force field was used for docking, at least 15 site points per reported site, a more limited definition of hydrophobicity, a fine grid, crop site maps at 4 from the nearest site point and reporting up to 5 sites were all necessary for this study.

### 3.6 Molecular Docking

All ligands were obtained directly from PubChem due to the size of the ligand library, we decided to make use of the QM Conformer and Tautomer Predictor [20] instead of using the conventional ligand preparation of Schrodinger [21]. This was done to ensure that we end up with quantum mechanical based minimum energy conformers prior to docking to the active site of the proteins of interest. The method generates the lowest energy tautomers or conformers for a set of structures with optional protonation or deprotonation. Due to the computational cost of this method, it is advised that this only be applied to small datasets.

### 3.7 Molecular Dynamics

Molecular dynamics (MD) simulations were conducted with the aid of Desmond [22]. The free protein structures as well as those complexed to Put, Spd, and Spn were prepared by using the TIP4P solvent model, 15 Å^3^ Orthorhombic boxes, OPLS4 force field, and each system was neutralized with an appropriate number of Na^+^ or Cl^-^ counterions (See Appendix 2). A 0.15 M NaCl solution was created to mimic physiological conditions. All MD simulations were done with the NPT ensemble at a temperature of 300 K and pressure of 1.01325 bar. Due to wall time limitations on the high-performance computer being used for the simulations, we had to run the MD in steps of 50 ns for HSP20, HSP40, and HSP60, 30 ns for HSP70, and 25 ns for HSP90. Upon completion of the simulations, the various intervals were combined into a single 200 ns trajectory for analysis.

## 4. Conclusions

Malaria is a major cause of death in many parts of the world, especially in Sub-Saharan Africa. The current treatment available in the market is not effective, due to the ever-changing parasite genome which then becomes resistant to the current drugs available in the market. This, therefore, needs urgent alternative treatment for malaria [24]. This study focused on three important polyamines namely: putrescine, spermidine, and spermine to establish the interaction with selected heat shock proteins. The interesting part we observed was that from this study was that all the focused polyamines were able to interact with selected heat shock proteins. But the interesting part shown by the MD was that the bigger the molecular chaperones the longer they interact with some polyamines. This interaction we observed between polyamines and heat shock proteins is a start toward the development of an alternative treatment for malaria. We will make follow-up studies based on the information we got from the current study.

## Author Contributions

All authors contributed equally to the study.

## Institutional Review Board Statement

Not applicable.

## Informed Consent Statement

Not applicable.

## Data Availability Statement

Data is contained within the article or supplementary material.

## Acknowledgments

The authors wish to acknowledge the University of Fort Hare, the National Research Foundation for student support, and the University of Cape Town for supporting Dr. S. Makumire. We would also like to acknowledge the Centre For High-Performance Computing, Rosebank, Cape Town for providing the computational resources used during this work.

